# Ecdysone signaling promotes expression of multifunctional RNA binding proteins essential for ovarian germline stem cell self-renewal in *Drosophila*

**DOI:** 10.1101/321109

**Authors:** Danielle S. Finger, Vivian V. Holt, Elizabeth T. Ables

## Abstract

Steroid hormones promote stem cell self-renewal in many tissues; however, the molecular mechanisms by which hormone signaling is integrated with niche-derived signals are largely uncharacterized. In the *Drosophila* ovary, the steroid hormone ecdysone promotes germline stem cell (GSC) self-renewal. Despite strong evidence that ecdysone modulates the reception of bone morphogenetic protein (BMP) signals in GSCs, transcriptional targets of ecdysone signaling that facilitate BMP reception are unknown. Here, we report that ecdysone signaling promotes the expression of the heterogeneous nuclear ribonucleoproteins (hnRNPs) *squid*, *hephaestus*, *Hrb27C*, and *Hrb87F* in GSCs. These hnRNPs functionally interact with ecdysone signaling to control GSC number and are cell autonomously required in GSCs for their maintenance. We demonstrate that hnRNPs promote GSC self-renewal by binding to transcripts essential for proper BMP signaling, including the BMP receptors *thickveins* and *punt*. Our findings support the model that stem cells coordinate local and long-range signals at the transcriptional and post-transcriptional levels to maintain self-renewal in response to physiological demand.

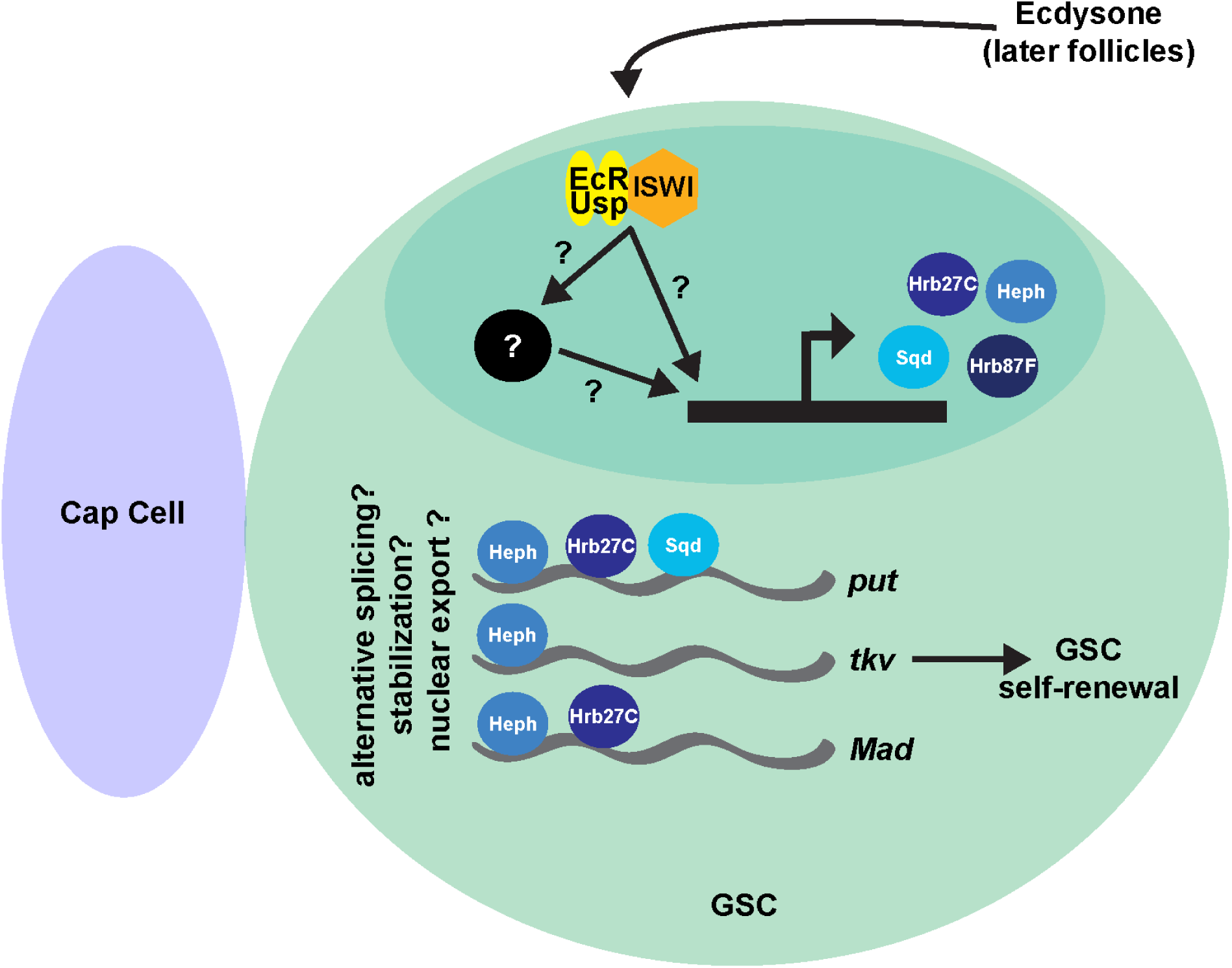
GRAPHICAL ABSTRACT. Ecdysone signaling regulates distinct hnRNPs that bind to BMP signaling targets to control GSC self-renewal.

**SUMMARY STATEMENT:** Ecdysone signaling promotes expression of heterogeneous ribonucleoproteins that modulate BMP-dependent germline stem cell self-renewal in the *Drosophila* ovary.

## INTRODUCTION

Stem cells are critical for tissue homeostasis and cellular diversity in developing and mature organs. Many stem cells divide asymmetrically, balancing long-term stem cell self-renewal with the production of progenitor cells that differentiate into functionally specialized cells (Chen et al., 2016; Ge and Fuchs, 2018; Gervais and Bardin, 2017). To ensure tissue integrity and proper organ function, stem cell self-renewal and proliferation must be tightly regulated and sensitive to changes in physiology over the lifetime of the organism. The molecular mechanisms connecting physiological signals to stem cell self-renewal, however, remain incompletely defined (Ables et al., 2012; Ghorbani and Naderi-Meshkin, 2016; Laws and Drummond-Barbosa, 2017). As stem cell decline contributes to age-related tissue degeneration (Keyes and Fuchs, 2018; Oh et al., 2014; Pan et al., 2007), understanding how physiological signals modulate stem cell activity may offer new strategies for optimization of tissue repair and regeneration *in vivo*.

Nuclear hormone receptors, which directly connect physiological signals to cellular responses, are well-suited to regulate stem cell self-renewal and proliferation. Nuclear hormone receptor ligands (e.g. steroid hormones) impact cell fate, proliferation, and survival in a wide variety of tissues. In the hematopoietic system, estrogen- and 27-hydroxycholesterol-induced activation of Estrogen Receptor α is essential for hematopoietic stem cell proliferation and subsequent expansion of erythropoiesis during pregnancy (Chapple et al., 2018; Nakada et al., 2014; Oguro et al., 2017). Mammary stem cells are likewise responsive to ovarian hormones (Asselin-Labat et al., 2010; Fu et al., 2017; Joshi et al., 2010). In the intestinal epithelium, excess circulating lipids induce expression of Peroxisome Proliferator-activated receptor δ, promoting intestinal stem cell self-renewal and predisposing cells to tumorigenesis (Beyaz and Yilmaz, 2016). Despite the therapeutic significance of nuclear hormone receptor signaling in tissue-resident stem cells, the molecular mechanisms by which these important receptors achieve tight regulation of stem cell activity remain largely uncharacterized (Ables and Drummond-Barbosa, 2017; Rafalski et al., 2012).

The *Drosophila melanogaster* ovary is a robust model system with which to elucidate the molecular mechanisms connecting local and systemic regulation of stem cell function. Ovaries are comprised of 14-16 ovarioles filled with maturing egg chambers or follicles, each of which will ultimately develop into a single egg (Spradling, 1993). Germline stem cells (GSCs) reside at the most anterior region of the ovariole (called the germarium; Fig. 1A-B) and are regulated by a complex network of paracrine and endocrine signaling mechanisms that maintain their self-renewal and proliferation (Xie, 2013). For example, GSCs are physically connected via adherens junctions to adjacent somatic cap cells (Fig. 1B), which secrete the bone morphogenetic protein (BMP) ligands Decapentaplegic (Dpp) and Glass bottom boat (Gbb) (Song et al., 2002; Xie and Spradling, 1998). Upon activation, BMP receptors Punt (Put) and Thickveins (Tkv) on GSCs suppress differentiation via activation of Mothers against decapentaplegic (Mad), which transcriptionally represses Bag of marbles (Bam), a primary differentiation factor (Chen and McKearin, 2003; McKearin and Ohlstein, 1995; Song et al., 2004). Translational control of differentiation factors is also critical for regulating GSC self-renewal and cystoblast differentiation (Slaidina and Lehmann, 2014). Asymmetric division of the GSC perpendicular to the cap cells produces another GSC and a cystoblast committed to differentiation. The cystoblast divides to form an interconnected 16-cell germline cyst. One cell of the cyst differentiates to form the oocyte, while the remaining cells become nurse cells, which ultimately load the oocyte with maternal factors essential for embryogenesis (Fig. 1B) (Spradling, 1993).

**Fig. 1.**
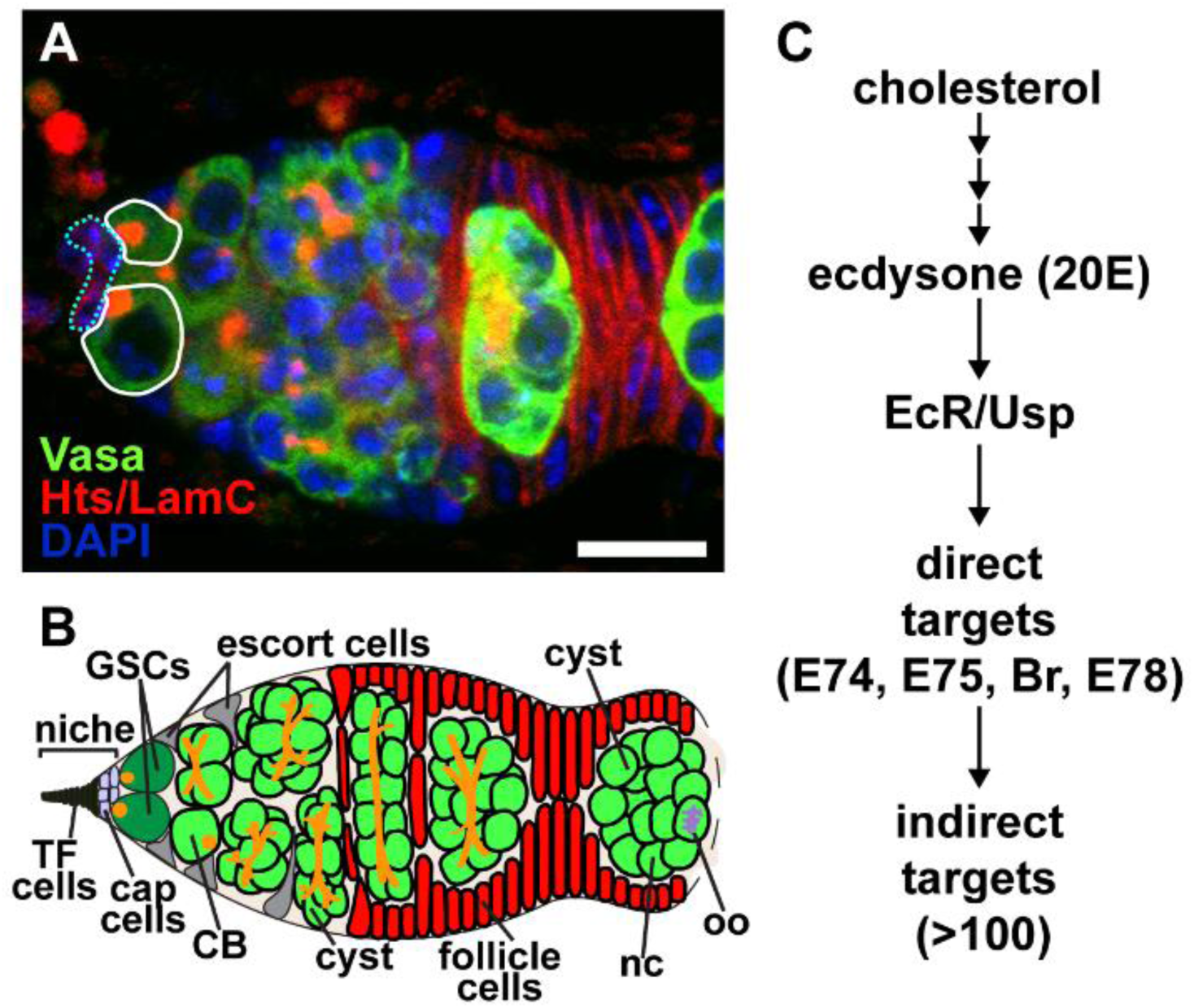
*Drosophila* oocyte development begins in the germarium. (A) *Drosophila* germarium labeled with anti-Vasa (green; germ cells) anti-Hts (red; fusomes and follicle cell membranes), and anti-LamC (red; nuclear envelope of cap cells). Solid lines demarcate GSCs, dotted blue lines demarcate cap cells. (B) GSCs (dark green) are anchored to a niche (composed of cap cells and terminal filament cells) at the anterior tip of each ovariole. Germ cells are characterized by the presence of a fusome (orange), which extends as germ cells divide. Escort cells (grey) signal to germ cells to promote differentiation. Follicle cells (red), surround the 16-cell germline cyst, giving rise to an egg chamber or follicle that leaves the germarium. (C) Diagram of the ecdysone signaling pathway depicting components with known roles in *Drosophila* oogenesis.

We and others have demonstrated that GSC self-renewal and proliferation are also modulated by physiological signals, including the steroid hormone ecdysone (Ables and Drummond-Barbosa, 2017; Belles and Piulachs, 2015). In adult females, ecdysone is produced in ovarian follicles (Uryu et al., 2015). Ecdysone signals, received by a heterodimer of nuclear hormone receptors Ecdysone Receptor (EcR) and Ultraspiracle (Usp), are necessary for GSC self-renewal and proliferation, germ cell differentiation, and follicle formation (Fig. 1C) (Ables and Drummond-Barbosa, 2010; Ables et al., 2016; Ameku and Niwa, 2016; Konig et al., 2011; Morris and Spradling, 2012). In GSCs, EcR/Usp signaling promotes reception of BMP signals, indicating that paracrine and endocrine signals are functionally integrated to regulate GSC behavior (Ables and Drummond-Barbosa, 2010). While this mechanism likely includes a functional interaction with the Iswi-containing chromatin regulator NURF, it remains unclear how EcR signaling modulates BMP signaling to maintain GSCs in a self-renewing fate.

In a recent genetic mosaic screen, we identified *Heterogeneous nuclear ribonucleoprotein at 27C* (*Hrb27C*), which encodes an RNA binding protein homologous to human *DAZ Associated Protein 1* (*DAZAP1*) and *Heterogeneous Nuclear Ribonucleoprotein A3* (*HNRNPA3*), as a putative target of ecdysone signaling that regulates GSC self-renewal (Ables et al., 2016). *Hrb27C* is a member of the heterogeneous nuclear ribonucleoproteins (hnRNPs) family of RNA binding proteins, whose critical functions include alternative splicing, stabilization of newly formed mRNA, transport in and out of the nucleus, and localization of mRNA (Chaudhury et al., 2010; Piccolo et al., 2014). For example, hnRNPs stabilize *E-cadherin* mRNAs to promote stem cell adhesion (Ji and Tulin, 2012), spatially restrict mRNAs in the *Drosophila* oocyte to establish embryonic axes, and mediate oligodendrocytic and neuronal mRNA trafficking in mice and humans (Piccolo et al., 2014). Further, evidence from mammals suggests that nuclear hormone receptor signaling can coordinately control transcription and pre-mRNA processing via recruitment of hnRNPs, perhaps as a mechanism to rapidly modify the cellular proteome in cells in response to physiological stimuli (Buoso et al., 2017; Curado et al., 2015; Dago et al., 2015; Zhou et al., 2015). Whether this mechanism controls stem cell self-renewal, however, has not been explored.

In this study, we investigate how ecdysone signaling might mechanistically regulate BMP signaling to control GSC self-renewal. We identify a subset of hnRNPs expressed in GSCs and dividing cysts, and demonstrate that expression of hnRNPs encoded by *Hrb27C*, *squid* (*sqd*; human homolog *HNRNPA/B*), *hephaestus* (*heph*; human homolog *Polypyrimidine Tract Binding protein 1*), and *Hrb87F* (human homolog *HNRNPA2/B*) are modulated by ecdysone signaling. We use spatially and temporally controlled loss-of-function analyses to show that *Hrb87F* is necessary to maintain the proper numbers of GSCs and that *Hrb27C*, *sqd*, and *heph* are intrinsically required in GSCs for their self-renewal. We show that Sqd and Heph bind RNAs encoding the BMP receptors Put and Tkv, and are essential for proper reception of BMP signaling. Taken together, our data support the hypothesis that ecdysone signaling promotes BMP signaling and GSC self-renewal in part by promoting the expression of specific hnRNPs. Our study offers new insights into how tissue-resident stem cells are modulated by the endocrine environment.

## RESULTS

### Many hnRNPs are expressed in GSCs and their differentiating daughters

Aberrant germline phenotypes, including dorsalized eggs and female sterility, were among the first biological processes attributed to mutations in *Drosophila* hnRNPs (Goodrich et al., 2004; Kelley, 1993; Matunis et al., 1994; Norvell et al., 1999). Yet while phenotypic reports of *Hrb27C* mutants hinted at potential function in the early germline (Yano et al., 2004), roles for hnRNPs in mitotically dividing germ cells have remained largely unexplored. We used available protein trap transgenes (Lye et al., 2014), reporters, and antibodies in wild-type ovaries to assess which of the 14 hnRNPs encoded in the *Drosophila* genome are expressed in the early germline (Fig. 2). Although reporters for most of the hnRNPs tested were expressed throughout the germline and soma (Fig. 2A-F), some distinct patterns emerged. *heph::GFP* (Besse et al., 2009) was expressed in GSCs and dividing cysts, but not in 16-cell cysts (Fig. 2G). *Hrb87F::GFP* (Singh and Lakhotia, 2012) was observed in nuclei of germ cells, but at much higher levels in somatic escort and follicle cells (Fig. 2H). Unlike the other hnRNPs tested, Syncrip (Syp) protein (McDermott et al., 2012) was exclusively observed in somatic cells in the germarium (Fig. 2I). Expression of hnRNPs in the germarium may indicate that, in addition to their roles in oocyte patterning in late oogenesis, hnRNPs are also important for the earliest stages of oocyte development.

**Fig. 2.**
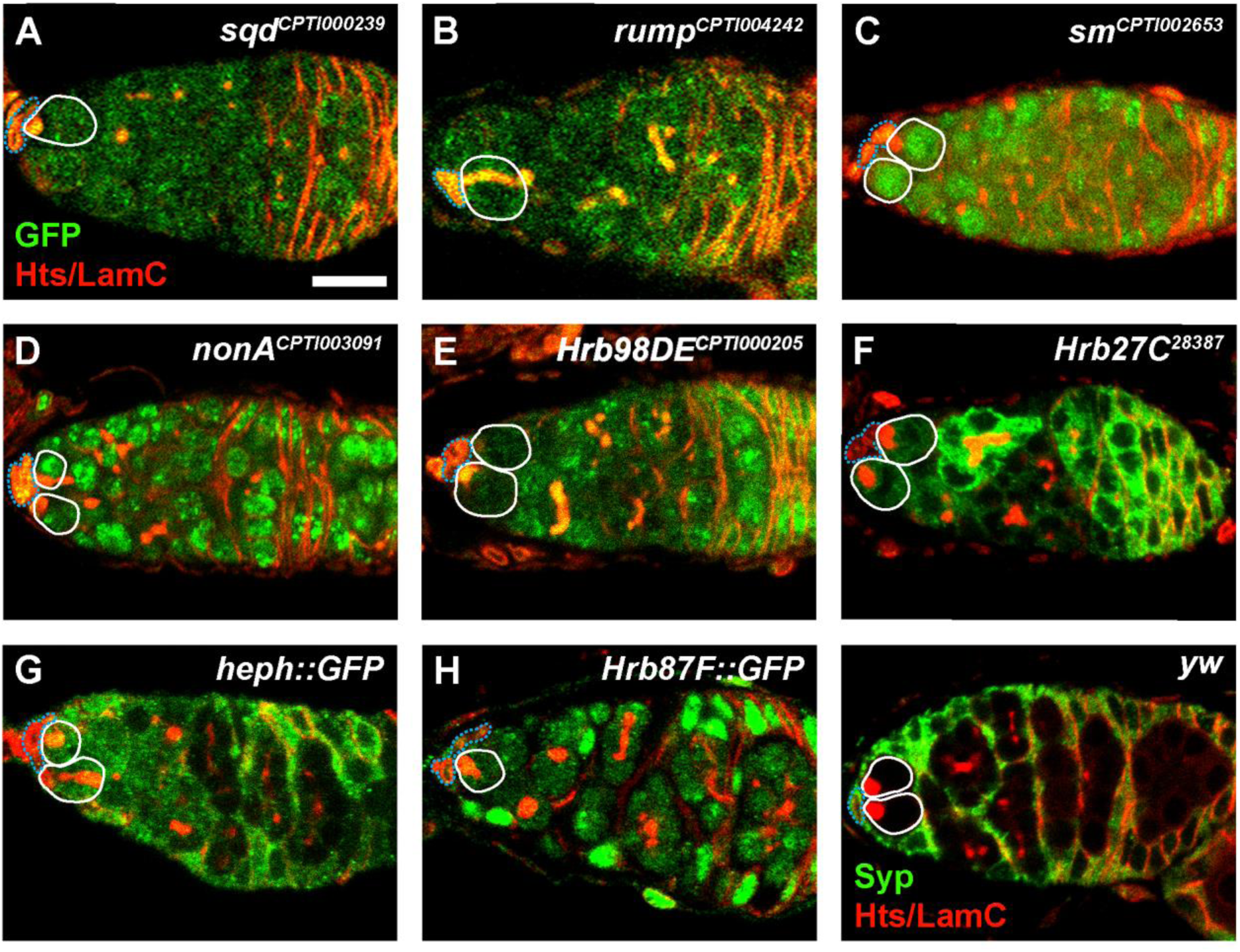
hnRNPs are expressed in distinct patterns in the germarium. (A-I) Representative germaria from GFP-tagged hnRNP transgenic flies labeled with anti-GFP (A-H) or wild-type flies labeled anti-Syncrip (I) and counterstained with anti-Hts (red; fusomes and follicle cell membranes), and anti-LamC (red; nuclear envelope of cap cells). Solid white lines demarcate GSCs, dotted blue lines demarcate cap cells. Scale bar = 10 µm.

### Ecdysone signaling promotes expression of hnRNPs in GSCs

We identified *Hrb27C* in a reverse genetic screen as a candidate target of ecdysone signaling in GSCs (Ables et al., 2016). Since hnRNPs are expressed in response to steroid hormones in other cell populations (Stoiber et al., 2016; Syed et al., 2017), and hnRNP complexes are found at many transcriptionally active, ecdysone-responsive chromosome regions in salivary gland polytene chromosomes (Amero et al., 1991; Amero et al., 1993), we asked whether other ovary-enriched hnRNPs are downstream of ecdysone signaling. We used quantitative reverse-transcription PCR (qRT-PCR) to measure selected hnRNP mRNA levels in whole ovaries from *EcR^ts^* mutant females, which have reduced ecdysone signaling (Ables and Drummond-Barbosa, 2010; Carney and Bender, 2000). We observed a statistically significant reduction in *Hrb27C, heph, sqd*, and *Hrb87F* levels in *EcR^ts^* mutant ovaries (Fig. 3A), suggesting that ecdysone signaling is required for proper expression of specific hnRNPs.

**Fig. 3.**
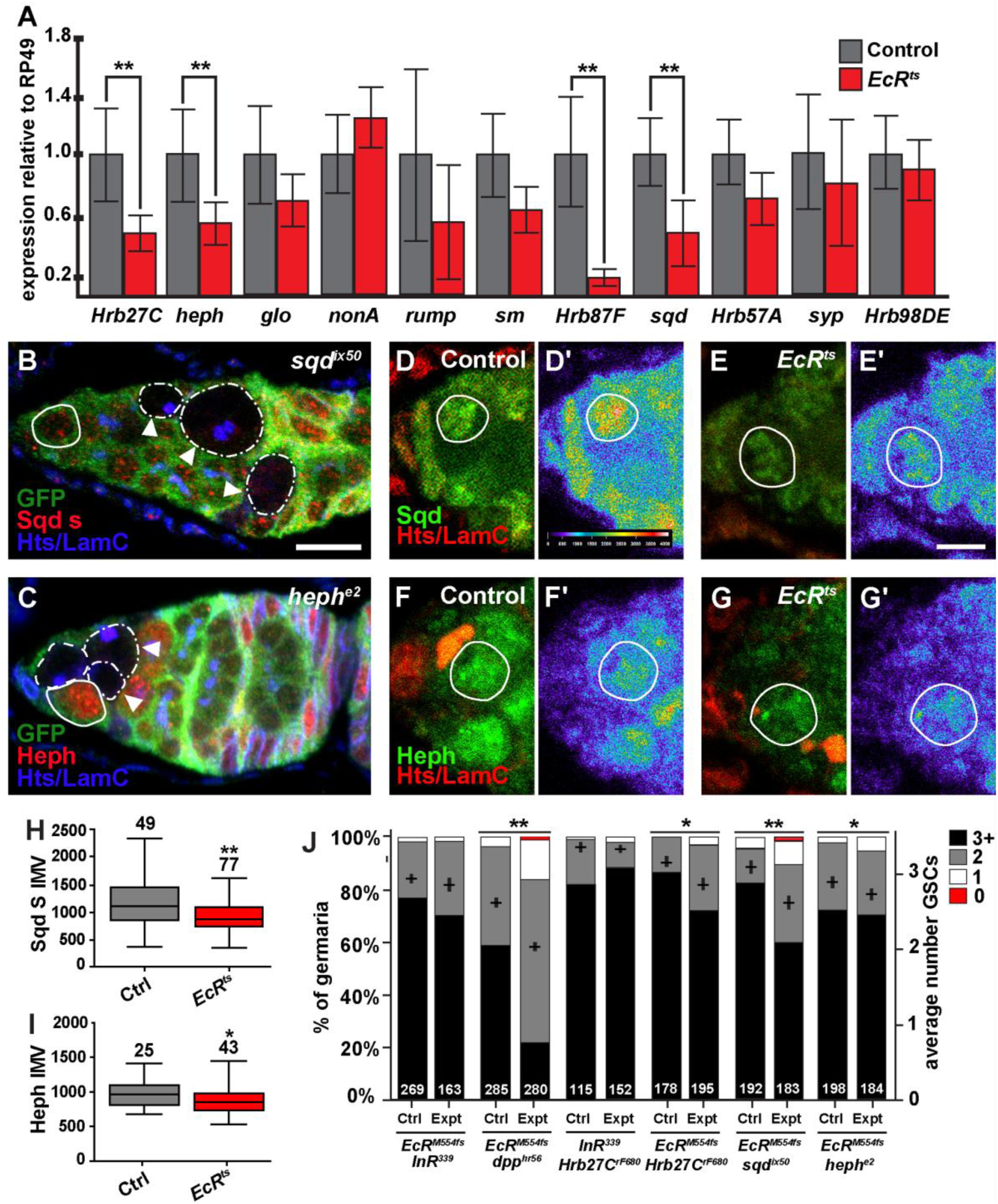
Ecdysone signaling transcriptionally regulates hnRNP expression in the ovary. (A) qRT-PCR of various hnRNPs in heterozygous sibling control (gray bars) and *EcR^ts^* (red bars) ovaries 3 days after temperature shift. Relative expression of three biological replicates normalized to *RP49* control. Error bars, mean ± SEM. (B) *sqd^ix50^* mosaic germaria stained with anti-GFP (green; wild type cells), anti-Hts (blue; fusomes, and follicle cell membranes), anti-LamC (blue; nuclear envelope of cap cells). (C) *heph^e1^* mosaic germaria stained with anti-GFP (green), anti-Hts (blue), anti-LamC (blue). GSCs are outlined in white (wild-type = solid line; mutant = dashed line). Irregular white line demarcates mutant cysts. Scale bar = 5 µm. (D, E) Expression of Sqd in sibling control (D) and *EcR^ts^* mutant (E) germaria counterstained with anti-Hts (red), anti-LamC (red). (F, G) Expression of Heph in sibling control (F) and *EcR^ts^* mutant (G) germaria counterstained with anti-Hts (red), anti-LamC (red). (D’-G’) Heat maps (purple < yellow < red) depicting Sqd (D’, E’) or Heph protein levels (F’, G’) in sibling control and *EcR^ts^* germaria. GSC nuclei are circled in white. Scale bar = 10 µm. (H, I) Fluorescence intensity mean value (IMV) of anti-Sqd (H) or Heph (I) antibody labeling. Number of GSCs analyzed is above bars. (J) Frequencies of germaria containing zero, one, two, or three or more GSCs (left y axis), and average number of GSCs per germarium (right y axis) 8 days after incubation at the restrictive temperature. The number of germaria analyzed is shown inside bars. Error bars, mean ± SEM. \*\**p* < 0.001; \**p* < 0.05.

Many transcripts, including hnRNPs, are produced by ovarian nurse cells and maternally deposited into the oocyte to support early embryonic development. We therefore asked whether decreased hnRNP mRNA in *EcR^ts^* ovaries resulted in a corresponding decrease in protein levels in the germarium. We obtained an anti-Sqd antibody that recognizes a Sqd isoform (Sqd s) expressed in early germ cells, but absent from *sqd^ix50^* mutant cells (Fig. 3B) (Norvell et al., 1999). Levels of Sqd were decreased throughout *EcR^ts^* germaria, including GSCs (Fig. 3D-E, H). Similarly, using an antibody specific for Heph (Fig. 3C), we observed reduced Heph protein in ecdysone-deficient GSCs (Fig. 3F-G, I). Taken together, these results demonstrate that ecdysone signaling promotes the expression of hnRNPs in GSCs.

We then sought to test whether hnRNPs might function together with ecdysone signaling to affect GSC maintenance. Ecdysone signaling regulates the transcription of hundreds of genes in a cell type-dependent manner (Stoiber et al., 2016), and hnRNPs function in combination to regulate RNA (Blanchette et al., 2009; Brooks et al., 2015; Huelga et al., 2012; McMahon et al., 2016). We reasoned that any evidence of genetic interaction between individual hnRNP loci and *EcR* would indicate that they function in a common pathway. We therefore performed a stringent genetic interaction assay to test whether heterozygous combinations of null alleles of *sqd*, *heph*, or *Hrb27C* and *EcR* would impact the number of GSCs per germarium, an indicator of normal GSC maintenance (Ables and Drummond-Barbosa, 2010). As a negative control, we tested double heterozygous females for *EcR^M554fs^* and the null *Insulin Receptor* (*InR^339^*) allele, because insulin-like peptides and ecdysone control GSC maintenance via distinct mechanisms (Ables and Drummond-Barbosa, 2010). As expected, we found no significant genetic interaction between the insulin and ecdysone pathways in controlling GSC number (Fig. 3J). Similarly, *Hrb27C^rF680^* and *InR^339^* did not genetically interact, arguing against the possibility that physiological signals *per se* function with hnRNPs to promote GSC maintenance. In contrast, double heterozygotes for *Hrb27C^rF680^* + / + *EcR^M554fs^*, *EcR^M554fs^*/+;*heph^e2^*/+, and *EcR^M554fs^*/+;*sqd^ix50^*/+ each had significantly fewer GSCs per germarium than single heterozygous controls (Fig. 3J). Intriguingly, we observed a stronger genetic interaction in *EcR^M554fs^* +/ + *dpp^hr56^* females than any of the hnRNP-containing double heterozygotes, indicating that one hnRNP locus alone is not sufficient to fully recapitulate the functional interaction between ecdysone signaling and BMP signaling in GSC maintenance (Ables and Drummond-Barbosa, 2010). Taken together, these results provide genetic evidence to support the model that *sqd*, *heph*, and *Hrb27C* function downstream of ecdysone signaling to regulate GSC maintenance.

### HnRNPs are independently and autonomously required in GSCs for their maintenance

Ecdysone signaling directly regulates GSC behavior; therefore, if hnRNPs are targets of ecdysone signaling, they should likewise be autonomously required in GSCs for their self-renewal. To test whether *Hrb27C*, *sqd*, and *heph* are essential for GSC self-renewal, we used *Flippase/Flippase Recognition Target* (*Flp/FRT*)-mediated mosaic recombination to inactivate their function in GSCs (Fig. 4). GSCs and their daughter cells carrying homozygous mutations in *sqd* (Fig. 4C), *heph* (Fig. 4D), or *Hrb27C* (Fig. 4E) were recognized by loss of GFP in mosaic germaria (Fig. 4A). Germaria were analyzed 8 days after clone induction, allowing negatively labeled cystoblasts/cysts to be cleared from the germaria and ensuring that GFP-negative cysts originated from a negatively labeled GSC. In most control “mock mosaic” germaria, where all cells are wild-type, GFP-negative GSCs were accompanied by GFP-negative cystoblasts/cysts, indicating that these GSCs self-renew and produce differentiating progeny (Fig. 4B, H). In contrast, a significant percentage of *sqd*, *heph*, and *Hrb27C* mutant mosaic germaria contained GFP-negative cysts without an accompanying GFP-negative GSC, indicating that the mutant GSC failed to be maintained in the niche (Fig. 4C-E, H).

**Fig. 4.**
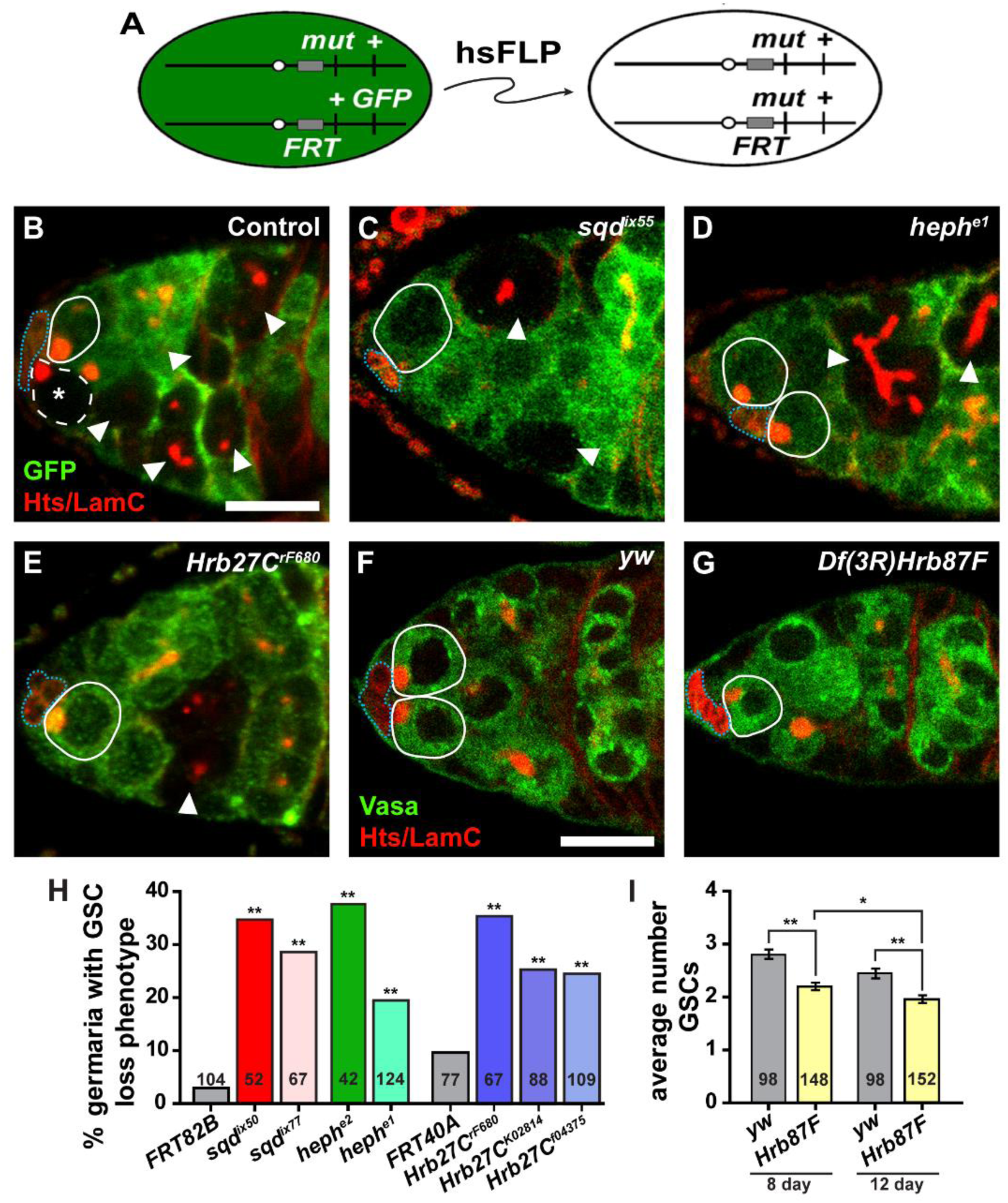
HnRNPs are required for GSC self-renewal. (A) The *FLP/FRT* technique was used to generate genetic mosaics. Mitotic recombination is mediated by heat-shock-induced expression of *flippase* (*hsFLP*). Homozygous mutant (mut) cells are identified by the absence of a GFP marker, which is linked to the wild-type allele. (B-E) Representative control (B), *sqd^ix50^* (C), *heph^e1^* (D), or *Hrb27C^rF680^* (E) mutant mosaic germaria labeled with anti-GFP (green; wild-type cells), anti-Hts (red; fusomes and follicle cells), and anti-LamC (red; nuclear envelope of cap cells). GSCs are outlined in white (wild-type = solid line; mutant = dashed line). Dotted blue line demarcates cap cells, arrowheads indicate GFP-negative cysts. (F-G) Sibling control (F) or *Df(3L)Hrb87F* mutant germaria labeled with anti-Vasa (green; germ cells), anti-Hts (red), and anti-LamC (red). GSCs are outlined. Scale bars = 5 µm. (H) Percentage of germline-mosaic germaria with a GSC loss event 8 days after clone induction. Numbers in bars represent the number of germline-mosaic germaria analyzed. (I) Average number of GSCs per germarium at 8 and 12 days after eclosion. \*\**p* < 0.001; \**p* < 0.05; Student’s two-tailed t-test. Error bars, mean ± SEM. Numbers in bars represent number of germaria analyzed.

Apparent loss of GSCs in our mosaic assay could be compounded by defects in cyst division or increased cyst death. To measure the rate of cyst production, we quantified cysts at each stage of mitotic division according to the morphology of the fusome, a specialized organelle that branches with each cyst division (de Cuevas and Spradling, 1998; Ong and Tan, 2010). The proportion of mutant and neighboring GFP-positive cystoblasts/cysts at each mitotic division was equivalent in *sqd^ix50^* and *Hrb27C^rF680^* mutant mosaic germaria (Fig. 5A). Using an antibody against cleaved Caspase-3 (Fig. 5B), we found no evidence of caspase-mediated apoptosis in *sqd^ix50^* (Fig. 5C), *heph^e2^* (Fig. 5D), or *Hrb27C^rF680^* (Fig. 5E) mutant mosaic germaria, suggesting that GSCs are not lost due to premature cell death. Further, although we observed *heph^e2^* mutant cysts at all stages of mitotic division (Fig. 4D, for example), *heph^e2^* mutant 4-, 8-, and 16-cell cysts were under-represented in mosaic germaria (Fig. 5A). These data suggest that *heph*, but not *sqd* or *Hrb27C*, is specifically necessary for cyst division, perhaps reflecting its prominent nuclear expression in early cysts (Fig. 2G and 3C).

**Fig. 5.**
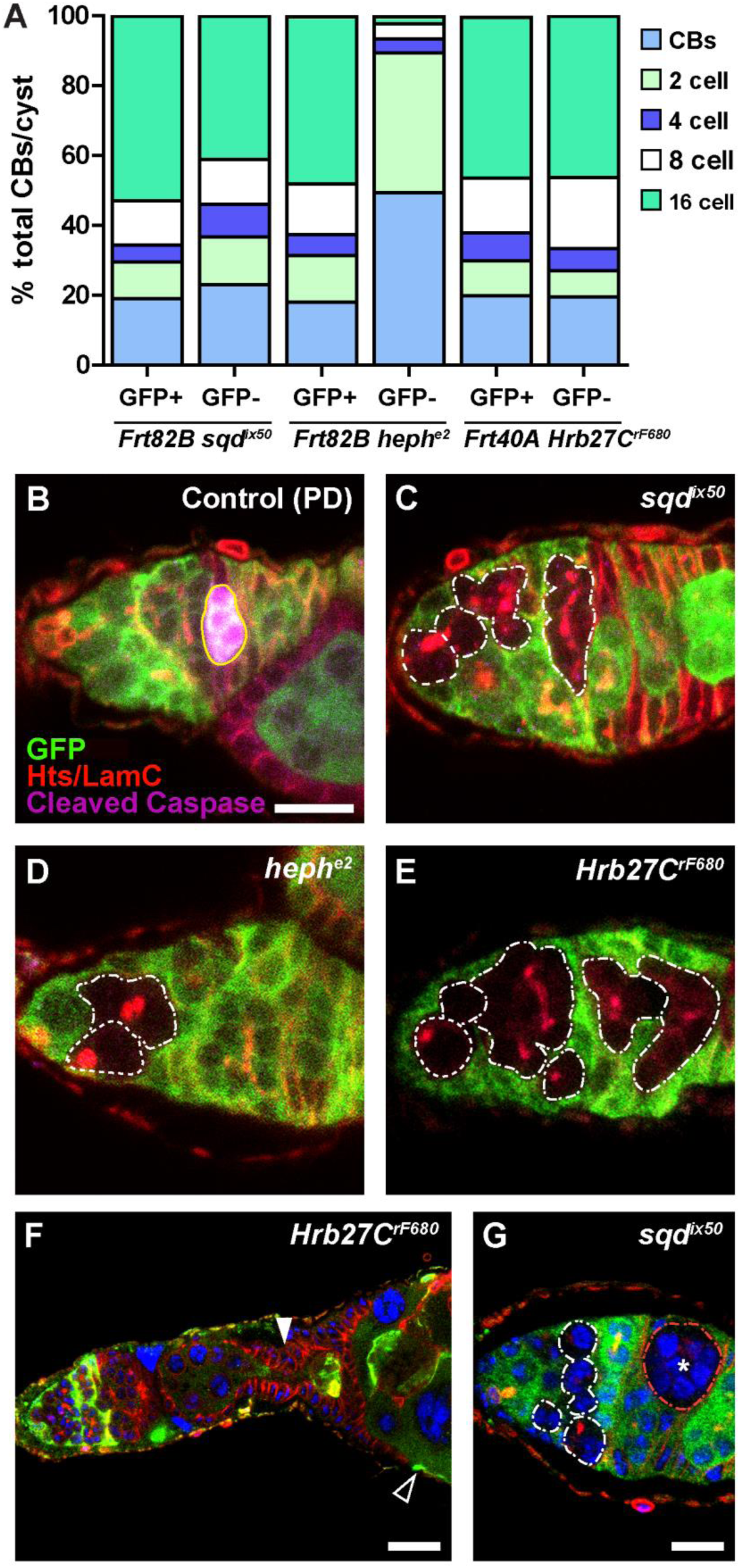
HnRNP mutant GSCs are not lost due to changes in the cell cycle. (A) Distribution of the average number of control (GFP-positive) or mutant (GFP-negative), GSCs, cbs, 2-, 4-, 8-, and 16-cell cysts per corresponding hnRNP mosaic germaria. Number germaria counted: sqd = 27, heph = 55, Hrb27C = 45. (B-E) Control (B), *sqd^ix50^* (C), *heph^e2^* (D), and *Hrb27C^rF680^* (E) mosaic germaria stained with anti-GFP (green; wild-type cells), anti-cleaved caspase (magenta, dying cyts), anti-Hts (red; fusomes and follicle cells), and anti-LamC (red; nuclear envelope of cap cells). GSCs are outlined in white (wild-type = solid line; mutant = dashed line), irregular white lines demarcate mutant cysts and yellow lines demarcate caspase positive cysts. Scale bars = 5 µm (F) *Hrb27C^rF680^* mutants have enlarged germaria. Mutant follicle cells fail to pinch off cyst (filled in arrowhead) and do not form a complete monolayer around cyst (outline arrowhead). (G) *sqd^ix50^* mutant display germline defects including circular cyst that lack a branching fusome (asterisk). Scale bars = 20 µm.

While *Hrb27C* mutant mosaics displayed no obvious early germline defects, we observed phenotypes outside of the germarium consistent with follicle formation defects, including dying cysts, fused cysts, and follicle cell overgrowth, indicating a critical role for *Hrb27C* in cyst encapsulation (Fig. 5F). Loss of *sqd* function from germline cysts also caused a variety of defects including rounded cysts lacking branched fusomes (Fig. 5G), suggesting that *sqd* serves several other roles in early cyst development and encapsulation.

Levels of *Hrb87F* mRNA were severely compromised in ecdysone-deficient ovaries (Fig. 3A). Although technical limitations prevented us from specifically inactivating *Hrb87F* function in GSCs using the *FLP/FRT* technique, a recent report demonstrating decreased female fecundity in *Df(3R)Hrb87F* mutants (Singh and Lakhotia, 2012) prompted us to investigate whether it is also necessary for GSC maintenance. Female homozygous *Df(3R)Hrb87F* mutants (see Experimental Procedures) survived to adulthood, but had multiple defects in oogenesis and were sterile (Singh and Lakhotia, 2012). Further, we observed fewer GSCs per germarium as compared to heterozygous sibling controls (Fig. 4F-G, I). Taken together, our data support the model that individual hnRNPs are independently necessary for GSC maintenance.

### *Hrb27C* is necessary for GSC proliferation

GSC self-renewal is, at least in part, dependent on efficient rates of GSC proliferation (Ables and Drummond-Barbosa, 2013). To test whether the inability of hnRNP mutant GSCs to be maintained reflects a reduced rate of proliferation, we measured the percentage of GSCs that incorporated the thymidine analog EdU in a 1 hour pulse in mosaic germaria. We did not observe statistically significant differences in rates of proliferation in *sqd^ix50^* (Fig. 6B, E) or *heph^e2^* (Fig. 6C, E) mutant GSCs, as compared to mock GFP-negative control GSCs (Fig. 6A, E). In contrast, significantly fewer *Hrb27C^rF680^* mutant GSCs incorporated EdU versus controls, indicating that *Hrb27C* specifically promotes GSC cell cycle progression. Taken together, these results argue against a general germ cell dysfunction due to RNA destabilization in hnRNP mutant cells, but instead support the model that *Hrb27C*, *heph*, and *sqd* control distinct cellular processes in germ cells.

**Fig. 6:**
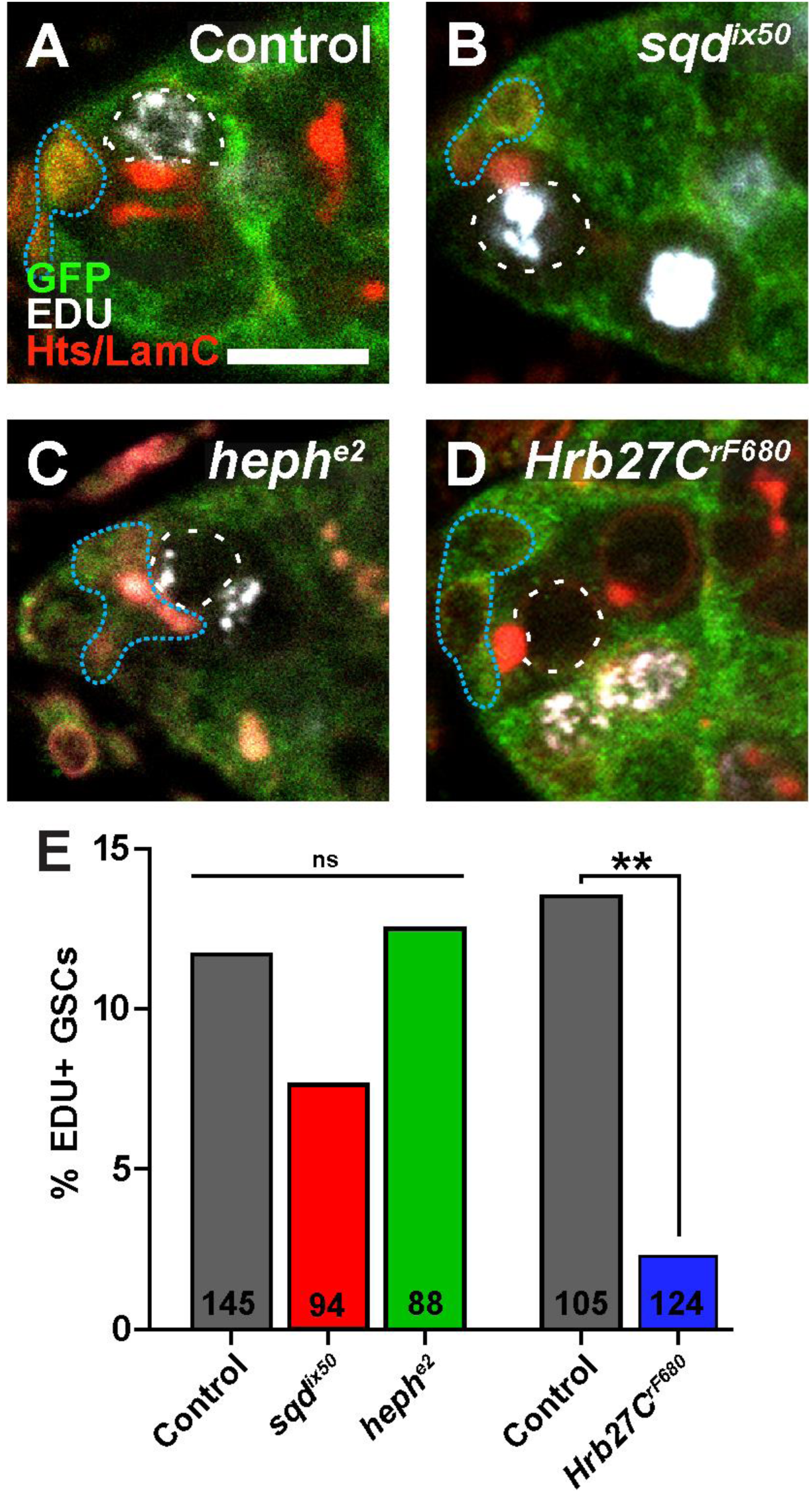
*Hrb27C* mutants display decreased GSC proliferation. (A-D) Control (A), *sqd^ix50^* (B), *heph^e2^* (C), and *Hrb27C^rF680^* (D) mosaic germaria stained with anti-GFP (green; wild-type cells), EdU (white; cells in S phase), anti-Hts (red; fusomes and follicle cells), and anti-LamC (red; nuclear envelope of cap cells). GSCs are outlined in white (wild-type = solid line; mutant = dashed line), dotted blue lines demarcate cap cells. Scale bars = 5 µm. **H**) Percentage of GFP-negative GSCs that are positive for EdU in mosaic germaria 8 days after heat shock. Numbers in the bars represent the number of GFP-negative GSCs analyzed. \*\**p* < 0.001.

### Loss of *heph*, *sqd*, or *Hrb27C* does not impair GSC adhesion to cap cells

HnRNPs frequently have overlapping targets (Blanchette et al., 2009; Brooks et al., 2015; Huelga et al., 2012; McMahon et al., 2016), at least in part because hnRNPs can form multi-protein complexes on RNA (Markovtsov et al., 2000). Previous studies indicated that *Hrb98DE* promotes GSC maintenance by binding the 5’ untranslated region of *E-cadherin*, promoting its translation and stabilizing the physical attachment of GSCs to the stem cell niche by adherens junctions (Ji and Tulin, 2012). E-cadherin localizes at the interface between cap cells and GSCs (Fig. 7) and is necessary for GSC self-renewal (Song et al., 2002). Because hnRNPs work in combination to promote mRNA splicing, transport, and stability (Matunis et al., 1992; Piccolo et al., 2014), we hypothesized that *Hrb98DE*, *sqd, heph*, and *Hrb27C* promote GSC self-renewal by binding a common target. To test whether loss of *Hrb27C*, *heph*, or *sqd* abrogated E-cadherin expression, we compared E-cadherin mean fluorescence intensity at the cap cell-GSC interface between hnRNP mutant GSCs and adjacent wild-type GSCs (Fig. 7). We found no significant difference in E-cadherin expression in *sqd^ix50^* (Fig. 7A-B), *heph^e2^* (Fig. 7C-D), or *Hrb27C^rF680^* mutant GSCs (Fig. 7E-F), indicating that impaired maintenance of *sqd, heph*, and *Hrb27C* mutant GSCs is not due to loss of a physical attachment to the niche.

**Fig. 7.**
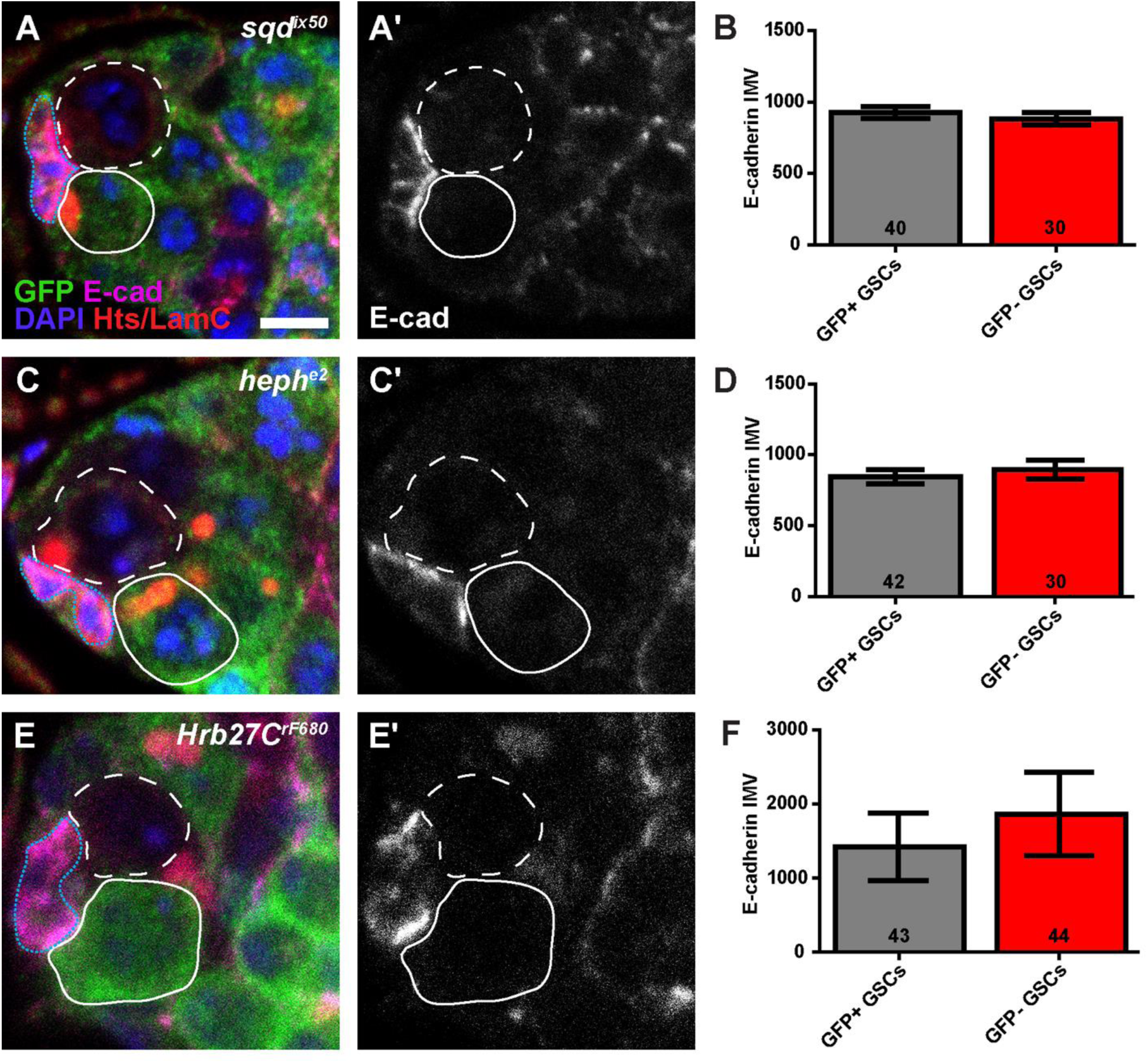
Loss of *Hrb27C*, *sqd*, or *heph* does not abrogate E-cadherin levels at the GSC/Cap cell interface. (A, C, E) *sqd^ix50^* (A), *heph^e2^* (C), and *Hrb27C^rF680^* (E) mosaic germaria labeled with anti-GFP (green, wild-type cells), anti-E-cadherin (magenta), anti-Hts (red; fusomes, and follicle cell membranes), anti-LamC (red; nuclear envelope of cap cells), and DAPI (blue; DNA). Greyscale images of E-cadherin alone are shown in A’, C’, and E’. GSCs are outlined in white (wild-type = solid line; mutant = dashed line), dotted blue line demarcates cap cells. (B, D, F). Fluorescence intensity mean value (IMV) of E-cadherin antibody labeling in adjacent control (wild-type; GFP+) and mutant (GFP-) GSCs. Error bars, mean ± SEM. Number of germaria analyzed within bars. Scale bar = 5 µm.

### *heph* and *sqd* promote BMP signaling in GSCs

Loss of ecdysone signaling leads to a failure in GSC self-renewal due to a decreased ability of GSCs to receive BMP signals (Ables and Drummond-Barbosa, 2010). Failure of *sqd, heph*, and *Hrb27C* mutant GSCs to self-renew is not due to reduced E-cadherin levels. We therefore asked whether premature differentiation contributed to the loss of *sqd, heph*, and *Hrb27C* mutant GSCs from the niche. BMP ligands, produced by somatic niche cells and received by GSCs, are essential for GSC self-renewal (Chen and McKearin, 2003; Song et al., 2004; Xie and Spradling, 1998). To test the ability of hnRNP mutant GSCs to respond to BMP signals, we measured the levels of phosphorylated Mothers against decapentaplegic (pMad), a well-characterized reporter of BMP pathway activation (Fig. 8A-E) (Kai and Spradling, 2003; Song et al., 2004). To control for technical variation in immunofluorescence staining, we calculated the ratio of nuclear pMad fluorescence intensity values in mutant (GFP-negative) GSCs and adjacent wild type (GFP-positive) GSCs in the same germarium (Fig. 8E). A GFP−/GFP+ ratio of 1 thus indicates equal levels of nuclear pMad in both cells. *heph^e2^* and *sqd^ix50^* mutant GSCs displayed significantly lower levels of nuclear pMad than neighboring wild-type GSCs (Fig. 6B-C, F), supporting the model that *heph* and *sqd* directly promote GSC self-renewal via BMP signaling. In contrast, we were unable to find a statistically significant difference in pMad fluorescence intensity in *Hrb27C^rF680^* mutant GSCs, suggesting that *Hrb27C* promotes GSC self-renewal predominately through another mechanism (perhaps via direct regulation of the GSC cell cycle; Fig. 6).

**Fig. 8.**
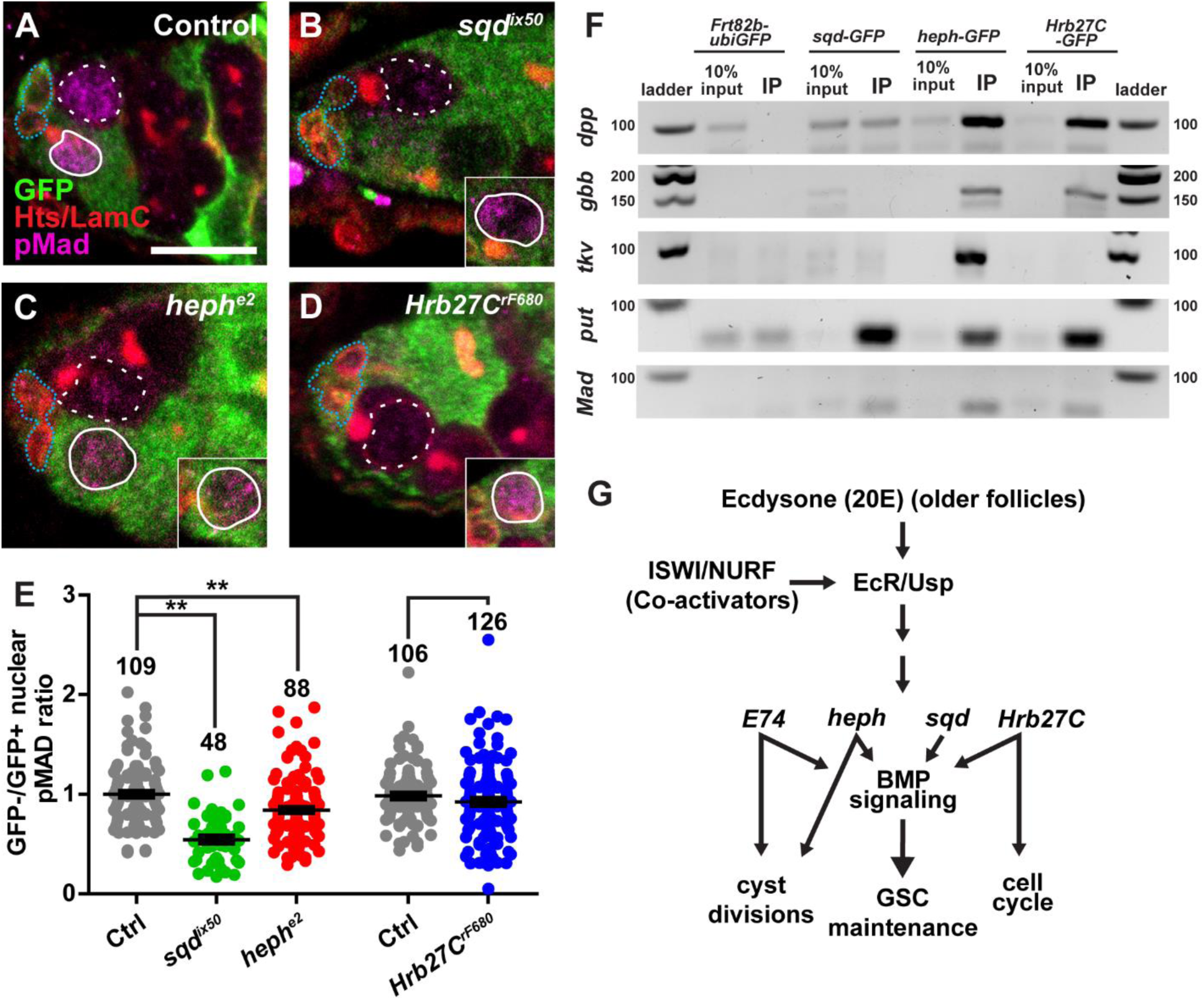
HnRNPs are necessary in GSCs to receive BMP signaling. (A-D) Control (A), *sqd^ix50^* (B), *heph^e2^* (C), and *Hrb27C^rF680^* (D) mosaic germaria labeled with anti-GFP (green, wild-type cells), anti-pMad (magenta), anti-Hts (red; fusomes, and follicle cell membranes), and anti-LamC (red; nuclear envelope of cap cells). Subset of WT GSC in same germarium. Scale bars = 5 µm. (E) Fluorescence intensity mean value (IMV) ratio of anti-pMAD antibody labeling in mutant GSCs /control GSCs within the same germarium. \*\**p* < 0.001; \**p* < 0.05; Student’s two-tailed t-test. Error bars, mean ± SEM. Number of germaria analyzed above bars. (F) RNA immunoprecipitation of GFP tagged hnRNPs with BMP signaling transcripts. (G) Proposed model of how ecdysone signaling regulates downstream targets that regulate the BMP signaling pathway to control GSC self-renewal.

We then asked how hnRNPs might regulate BMP signaling. We performed RNA immunoprecipitation followed by RT-PCR in ovarian extracts to test whether GFP-tagged Sqd, Heph, or Hrb27C were in a ribonucleoprotein complex with transcripts essential for proper BMP signaling (Fig. 8F). We first tested whether the hnRNPs could bind transcripts encoding the BMP ligands *dpp* or *gbb*, which are essential for GSC maintenance (Song et al., 2004). Interestingly, Heph and Hrb27C, but not Sqd, immunoprecipitated with both ligands. We then tested whether RNAs encoding the BMP receptors *tkv* and *put* or the BMP-regulated transcription factor *Mad* were in complex with the hnRNPs. While all three hnRNPs efficiently bound *put* and, to a lesser extent, *Mad*, only Heph also bound *tkv*. Taken together with our functional analysis of *heph* and *sqd* mutant GSCs, these results support the model (Fig. 8G) that Heph and Sqd autonomously promote BMP signaling primarily by binding to RNAs encoding BMP receptors, perhaps to regulate their stability, splicing, or intracellular transport.

## DISCUSSION

Germline stem cell self-renewal is regulated by a combination of paracrine and hormonal signals; however, the molecular mechanisms by which these critical signals converge in GSCs has remained largely unclear. In this study, we demonstrate for the first time that distinct hnRNPs are transcriptionally regulated by the steroid hormone ecdysone in the *Drosophila* ovary. We also show that conserved hnRNPs *sqd* (*hnRNPA/B*), *heph* (*PTBP1*), *Hrb87F* (*hnRNPA2/B*), and *Hrb27C* (*DAZAP1/hnRNPA3*) are essential for GSC self-renewal. Our genetic and biochemical analyses suggest that Heph, Sqd, and Hrb27C promote GSC self-renewal via distinct molecular mechanisms, at least in part via regulation of RNA necessary to sustain proper BMP signaling. Taken together, our results demonstrate one possible molecular mechanism by which two signaling pathways might work together to regulate stem cell activity in response to physiological demand.

### hnRNPs are necessary for early germ cell development

Gene expression involves the formation of distinct RNP complexes containing nascent transcripts, hnRNPs, and translation initiation factors (Bjork and Wieslander, 2017). Post-transcriptional regulation, translational repression, and the formation of RNP complexes are hallmarks of germ cell development, and are critical for meiosis in a variety of organisms (Jin and Neiman, 2016; Licatalosi, 2016; Nousch and Eckmann, 2013; Percipalle, 2014; Slaidina and Lehmann, 2014). Indeed, one of the first biological roles attributed to hnRNPs in *Drosophila* oogenesis was the establishment of anterior-posterior and dorsoventral axes in oocytes (Kelley, 1993; Matunis et al., 1994). In particular, Sqd, Hrb27C, and Glo are necessary for translational repression and localization of *gurken*, *nanos*, and *oskar* mRNAs, whose spatially-regulated translation establishes concentrated areas of asymmetrically distributed protein in the oocyte (Goodrich et al., 2004; Huynh et al., 2004; Kalifa et al., 2009; Kalifa et al., 2006). More recently, *Hrb98DE*, *Hrb27C*, and *Protein on ecdysone puffs* (*Pep*) were identified in genetic screens for novel regulators of GSC self-renewal (Ables et al., 2016; Yan et al., 2014). It should perhaps come as no surprise then that hnRNPs are expressed in and essential for GSC function and germ cell development. What is surprising, however, is that even though many of the hnRNP-dependent factors that promote oocyte axes establishment (e.g. *nanos*, *vasa*) are also required for GSC self-renewal (Slaidina and Lehmann, 2014), we found no evidence that *sqd*, *Hrb27C*, or *heph* modulate Nanos or Vasa protein levels in GSCs or cystoblasts (data not shown). Taken together, our data suggests a broad role for hnRNPs in the regulation of key events in oocyte development.

HnRNPs are a diverse family of RNA binding proteins that regulate post-transcriptional processing, maturation, and nuclear export of RNA polymerase II-dependent transcripts (Levengood and Tolbert, 2018). Attempts to identify transcripts bound by hnRNPs in *Drosophila* have yielded thousands of putative targets and illuminated extensive cross-regulatory interactions (Blanchette et al., 2009; Brooks et al., 2015; McMahon et al., 2016; Stoiber et al., 2015). Our data suggest that Sqd, Heph, and Hrb27C post-transcriptionally promote the expression of key components of the BMP signaling pathway in GSCs. In support of this model, recent studies profiling RNA targets of Hrb27C in *Drosophila* neuronal cell lines identified several transcripts essential for BMP signaling (McMahon et al., 2016; Xu et al., 2018). Given the complexity of the *sax*, *tkv*, and *put* gene loci, we postulate that hnRNPs regulate the splicing or stability of these essential BMP signaling transcripts in GSCs. We cannot exclude the possibility, however, that the hnRNPs regulate other as-yet-unidentified transcripts critical for promoting GSC self-renewal. Future experiments will be necessary to elucidate the complex molecular mechanisms underlying these effects.

### The intersection between hormone signaling and translational control: a common mechanism promoting cell fate?

The cellular response to ecdysone signaling is complex, exhibiting both spatial and temporal specificity (Yamanaka et al., 2013). Ecdysone-dependent expression of hnRNPs, however, does not appear to be specific to ovarian cells (Beckstead et al., 2005; Gauhar et al., 2009; Stoiber et al., 2016; Syed et al., 2017). In the *Drosophila* central nervous system, ecdysone signaling temporally promotes expression of Syp to coordinate neuroblast terminal differentiation with organismal development (Doe, 2017; Syed et al., 2017). Further, a recent study profiling the transcriptome in *Drosophila* cell lines demonstrated that hnRNP expression is frequently induced in response to ecdysone (Stoiber et al., 2016). We speculate that hormone signaling coordinates chromatin state, transcriptional regulation, and post-transcriptional processing to broadly regulate gene expression.

We previously demonstrated that ecdysone signaling functionally interacts with the Iswi-containing nucleosome remodeler NURF in GSCs to promote their self-renewal (Ables and Drummond-Barbosa, 2010). *Iswi* mutants phenocopy loss-of-function of the ecdysone response gene *E74*; specifically, both mutant *E74* and *Iswi* GSCs fail to be maintained over time, cycle more slowly, and exhibit decreased levels of pMad, suggesting that they fail to receive or properly interpret BMP signals (Ables and Drummond-Barbosa, 2010; Xi and Xie, 2005). While *E74* and *usp* mutant GSCs resulted in reduced levels of Iswi protein, reductions in ecdysone signaling did not impair *Iswi* or *E(bx)* transcription (Ables and Drummond-Barbosa, 2010). Since NURF binds EcR in vitro and is essential for transcription of EcR-dependent transcripts (Ables and Drummond-Barbosa, 2010; Badenhorst et al., 2005; Kugler et al., 2011), we favor the model that NURF functions as an EcR co-activator (see Graphical Abstract). Although *E74* is clearly required for GSC self-renewal and germ cell mitotic divisions, it is unlikely that it is the only ecdysone-response gene that is essential in GSCs and their daughters. Indeed, ecdysone signaling is well-known to regulate hundreds of transcriptional targets, both directly and indirectly via activation of early response gene-mediated transcriptional cascades. In our current work, we identify a set of hnRNPs that are transcriptionally regulated downstream of EcR. Although the phenotypes of *heph*, *sqd*, and *Hrb27C* mutants are individually relatively mild, we speculate that the combination of these ecdysone-sensitive targets into distinct functional complexes helps fine-tune the cellular response to changes in hormone levels in GSCs.

We envision three possible models by which ecdysone may coordinate unique hnRNP-containing RNP complexes in germ cells to regulate differentiation. First, ecdysone signaling via EcR/Usp may transcriptionally promote hnRNP expression in specific cells or at specific stages of differentiation. Increased receptor concentration, co-activator or co-repressor activity, or EcR isoform specificity could bias RNP complex formation by up-regulating specific hnRNPs in specific cells. Second, ecdysone signaling could modify the chromatin landscape in specific cell lineages or stages of differentiation. EcR/NURF activity could promote open regions of chromatin at hnRNP gene loci, such that other lineage-specific transcription factors have access promote transcription. Lastly, ecdysone signaling could be positively reinforced in specific cell types by the post-transcriptional regulation of early response genes by hnRNPs. Unique hnRNP complexes have been observed at sites of ecdysone-dependent transcription, including the *E74* and *E75* loci (Amero et al., 1993). Taken together, this presents an attractive model by which steroid hormone signals are uniquely interpreted by cells at different stages of development or differentiation, or in specific lineages within a given tissue. Given the recent interest in hnRNP regulation as a causative factor in neurodegenerative disorders (Geuens et al., 2016), our study may provide novel insight into the origins of complex human diseases.

## MATERIALS AND METHODS

### *Drosophila* strains and culture conditions

Flies were maintained at 22°-25°C on a standard medium containing cornmeal, molasses, yeast and agar (Nutrifly MF; Genesee Scientific) supplemented with yeast. For all experiments, unless otherwise noted, flies were collected 2 to 3 days after eclosion and maintained on standard media at 25°C. Flies were supplemented with wet yeast paste (nutrient-rich diet) 3 days before ovary dissection. Genes/alleles with multiple names are referenced using FlyBase nomenclature (www.flybase.org) for simplicity.

The following alleles were used for protein expression and RNA immunoprecipitation: Sqd^*CPTI000239*^, rump^*CPTI004242*^, sm^*CPTI002653*^, nonA^*CPTI00309*^, and, Hrb98DE^*CPTI000205*^, (Kyoto) (Lowe et al., 2014), *PTB::GFP* (referred to as Heph::GFP) (Besse et al., 2009), *Hrb87F::GFP* (Singh and Lakhotia, 2015), and *Hrb27C^28387^* (Sarov et al., 2016).

For genetic interaction analyses, the following alleles were used: *Hrb27C^rF680^*(Goodrich et al., 2004)*, Hrb27C^02647^* (Hammond et al., 1997), *InR^339^* (LaFever and Drummond-Barbosa, 2005). *dpp^hr56^* (Bloomington #36528) (Xie and Spradling, 1998), *tkv^7^* (Bloomington #3242) (Terracol and Lengyel, 1994), *EcR^M554fs^* (Bloomington #4894) (Carney and Bender, 2000), *E74^neo24^* (Bloomington # 10262) (Fletcher et al., 1995), *E74^DL-1^* (Bloomington #4435) (Fletcher et al., 1995), *heph^e2^* (Dansereau et al., 2002), and *sqd^ix50^* (Kelley, 1993). Average stem cell number of double heterozygote mutants were compared to average stem cell number of controls (single heterozygotes carrying a balancer). Data was analyzed using Student’s t-test.

*EcR*-deficient ovaries (referred to as *EcR^ts^*) were created using temperature-sensitive *EcR^A438T^* mutants in trans with *EcR^M554s^* (Ables and Drummond-Barbosa, 2010; Carney and Bender, 2000) null mutants. These flies were raised at a permissive temperature (18°C) until eclosion, then incubated at the restrictive temperature (29°C) for 3 days and supplemented with wet yeast paste prior to dissection (Ables and Drummond-Barbosa, 2010).

### Genetic mosaic generation and stem cell analyses

Genetic mosaic analysis via *flippase/flippase recognition target* (*Flp/FRT*) (Xu and Rubin, 2012) used the following alleles on *FRT*-containing chromosomes: *sqd^ix50^* and *sqd^ix77^(Kelley, 1993)*, *heph^e1^(Dansereau et al., 2002)*, *heph^e2^*, *Hrb27C^rF680^*, *Hrb27C^K02814^* (Kyoto #111072), *Hrb27C^f04375^* (Kyoto #114656) (Spradling et al., 1999). Other genetic tools are described in FlyBase. Genetic mosaics were generated using *FLP/FRT*-mediated recombination in 1-3 day old females carrying a mutant allele *in trans* to a wild-type allele (linked to a *Ubi-GFP* or *NLS-RFP* marker) on homologous *FRT* arms with a *hs-FLP* transgene, as previously described (Hinnant et al., 2017; Laws and Drummond-Barbosa, 2015). Flies were heat shocked at 37°C twice per day 6-8 hours apart for 3 days, then incubated at 25°C on standard media supplemented first with dry yeast, then with wet yeast 3 days prior to dissection. Flies were dissected 8 days after clone induction. Wild-type alleles (*FRT40A* or *FRT82B*) were used for control mosaics. GSCs were identified by the location of their fusomes adjacent to the cap cells (de Cuevas and Spradling, 1998). GSC loss was measured by the number of germaria that contain a GFP-negative cyst (generated from the original GFP-negative stem cell) but lack a GFP-negative GSC, compared to the total number of germaria containing a germline clone (Laws and Drummond-Barbosa, 2015). Results were analyzed by Chi-square tests using Microsoft Excel. To measure stem cell loss in *P{w+Tsr+}/P{w+Tsr+}; ry Df(3R)Hrb87F/ry Df(3R)Hrb87F* (referred to as *Df(3R)Hrb87F*) (Singh and Lakhotia, 2012), flies were raised at 25°C and dissected 8 and 12 days after eclosion. GSC loss was measured by the average number of GSCs per germarium in mutants compared to heterozygous sibling controls.

### Quantitative RT-PCR

*EcR^ts^* mutants (see above) were raised at a permissive temperature (18°C) until eclosion, then incubated at the restrictive temperature (29°C) for 3 days (Ables and Drummond-Barbosa, 2010). Ovaries were dissected in RNAlater (Ambion) and stored at −20°C. Samples were comprised of 10 whole ovary pairs. Total RNA was extracted using an RNAqueous Total RNA isolation kit (Thermo Fisher). RNA was treated with a Turbo DNA-free kit (Thermo Fisher) following manufacturer’s instructions. RNA quality was tested using agarose gel electrophoresis and concentration was quantified using a Nanodrop Lite spectrophotometer (Thermo Fisher). Complementary DNA was created by reverse transcription using an iScript cDNA kit (Bio-Rad) following manufacturer’s instructions and using 500 ng of input RNA for each sample. Quantitative PCR was performed using a CFX96 Touch Real-Time PCR detection system (Bio-Rad). Primers (FlyPrimerBank) (Hu et al., 2013) amplified a region common to all predicted transcripts from the locus of interest: *Hrb27C* (F 5’-GGAAGACGAGAGGGGCAAAC-3’ R 5’-GAAGTAGCGCGACAGGTTCT-3’), *heph* (F 5’-ACCGCCCATAGCGACTACA-3’ R 5’-TTGAGCTGTTTGCATTGTTGC-3’), *glo* (F 5’-AACGCAGACGTGCAATTTAAC-3’), *nonA* (F 5’-GCCCAGAATCAAAACCAGAACC-3’ R 5’-CGAACCCACCCTTGTTGTTTC-3’), *rump* (F 5’-GGACGCTAGTAACTCGGTGG-3’ R 5’-CTTCAGATCCTGCCAACGGT-3’), *sm* (F 5’-TGGTGCAAATGGGAGACGC-3’ R 5’-AAGCGATCTGTATCTTGCCAC-3’), *Hrb87F* (F 5’-CACGTACTCCCAGTCGTACAT-3’ R 5’-GCAGGCACTCTTCATCGTGA-3’), *sqd* (F 5’-CGCAAAGGATTCTGCTTCATCA-3’ R 5’-CACGCTTAACATCGACCTCC-3’), *syp* (F 5’-CTCTCTAGCCAAACCCCC-3’ R 5’-ACGAGCACGCAGAATCTCC-3’), *Hrb57A* (F 5’-ATGAACTTTGACCGCGTATATGC-3’ R 5’-CTCCGTTACGATTGTTTCCCC-3’), *Hrb98DE* (F 5-TGGCTTGGACTACCGTACCA R 5’-TGGGAATAGGTGATGAATCCGAA-3’). Each analysis was performed in triplicate using iQ SYBR Green Supermix kit (Bio-Rad). Samples were standardized to a *RP49* control. Quantification was performed using the Bio-Rad CFX Manager program.

### RNA immunoprecipitation

3-6 day old *hnRNP-GFP* tagged females (see above), fed for 3 days with yeast paste, were dissected in DEPC-treated PBS. *hsFLP;FRT82B,ubiGFP* females served as controls. Four biological replicates using 50 pairs of ovaries per sample were dissected for each genotype. Immunoprecipitation and RNA extractions were done following manufactures instructions using the magna-RIP kit (Millipore-Signa) and rabbit anti-GFP (ab6556, Abcam, 1:2000). cDNA was created as described above. 10% of input RNA was used as a control. Samples were amplified by PCR for 40 cycles with iQ SYBR Green Supermix kit and PCR products separated on a 3% agarose gel and imaged using ethidium bromide. Primers (FlyPrimerBank) (Hu et al., 2013) amplified a region common to all predicted transcripts from the locus of interest: *dpp* (F 5’-TGGCGACTTTTCAAACGATTGT-3’ R 5-CAGCGGAATATGAGCGGCAA-3’), *gbb* (F 5’-GAGTGGCTGGTCAAGTCGAA-3’ R 5’-GAAGCCGATCATGAAGGGCT-3’), *tkv* (F 5’-ATGGAACCTGCGAGACCAGAC-3’ R 5’-CTCCTCGTACATCCCGGT-3’), *put* (F 5’-TGTGAAACACGGATAGAGCAC-3’ R 5’-GTTGACCGACCAAAGGACATAG-3’), *Mad* (F 5’-CCGAATGCCTATCCGACTCC-3’ R 5’-ATCCGTGGTGGTAGTTGCAG-3’).

### Immunofluorescence and microscopy

Ovaries were dissected, fixed, washed, and blocked as previously described (Ables et al., 2016; Hinnant et al., 2017). Briefly, ovaries were dissected and teased apart in Grace’s media (Lonza or Caisson Labs) and fixed using 5.3% formaldehyde in Grace’s media at room temperature for 13 minutes. Ovaries were washed extensively in phosphate-buffered saline (PBS, pH 7.4; Thermo Fisher) with 0.1% Triton X-100, then blocked for three hours in a blocking solution consisting of 5% bovine serum albumin (Sigma), 5% normal goat serum (MP Biomedicals) and 0.1% Triton-X-100 in PBS. The following primary antibodies were diluted in block and used overnight at 4°C: mouse anti-Lamin C (LamC) [LC28.26, Developmental Studies Hybridoma Bank (DSHB), 1:100], mouse anti-Hts (1B1, DSHB, 1:10), rabbit anti-GFP (ab6556, Abcam, 1:2000), chicken anti-GFP (ab13970, Abcam, 1:2000), guinea pig anti-Syncrip (a gift from I. Davis, 1:5000) (McDermott et al., 2012), rat anti-E-cadherin (DCAD2, DSHB, 1:20), rabbit anti-pMad [(Smad3) phospho S423 + S425, ab52903, Abcam/Epitomics, 1:50], rat anti-Heph (PTB, a gift from A. Ephrussi, 1:1000) (Besse et al., 2009), mouse anti-Sqd (Sqd S 1B11, DSHB, 1:7). Samples incubated with pMad were permeabilized with 0.5% Triton-X100 in PBS for thirty minutes before blocking. Samples were incubated with Alexa Fluor 488-, 568- or 633-conjugated goat-species specific secondary antibodies (Molecular Probes; 1:200) and counterstained with DAPI (Sigma 1:1000 in PBS). Ovaries were then mounted in 90% glycerol containing 20.0 µg/mL N-propyl gallate (Sigma). Data was collected using a Zeiss LSM 700 laser scanning confocal microscope. Images were analyzed using Zen Blue 2012 software and images were minimally and equally enhanced via histogram using Zen and Adobe Photoshop CS6.

### Quantification of fluorescence intensity in GSCs

Fluorescence intensity in confocal sections was measured via ZEN Blue 2012 (Zeiss) by manually demarcating individual GSC and measuring nuclear intensity mean values (IMV; gray value/pixel) at the z-level containing the largest nuclear diameter for the specific antibody analyzed. Because of slight variations in pixel intensity among stain sets, IMVs for each fluorescent protein were calculated for a minimum of 30 individual GSCs. Controls are adjacent GFP-positive (wild type) GSCs within the same germarium. To normalize pMAD quantification, IMVs were evaluated as a ratio of GFP−/GFP+ GSCs within the same germarium, where a ratio equal to 1 represented equivalent intensity in both cells. Statistical analysis was performed using Student’s t-test, in either Prism (GraphPad) or Excel.

## AUTHOR CONTRIBUTIONS

D.S.F. and E.T.A. conceived and designed experiments, analyzed data, interpreted results, and wrote the manuscript. D.S.F. performed the experiments and managed the project. V.V.H. assisted with experiments and data collection (Figure 1 and Table 1). All authors provided critical feedback and helped shape the research, analysis, and manuscript.

## ACKNOWLEDGMENTS

Many thanks to: D. Drummond-Barbosa, A. Norvell, T. Schüpbach, W. Brook, I. Davis, D. Ennis, A. Ephrussi, S. Lakhotia, P. Lasko, the Bloomington and Kyoto Stock Centers, and the Developmental Studies Hybridoma Bank for fly stocks and antibodies; the ECU Department of Biology Genomics and Microscopy Core Facilities; and A. Norvell, B. Thompson, C. Geyer, J.-L. Scemama, L. Weaver, members of the Ables laboratory, and anonymous reviewers for helpful discussions and critical reading of this manuscript.

